# Heparin-modified aligned collagen scaffolds enhance *in vitro* myogenesis

**DOI:** 10.1101/2025.10.15.681268

**Authors:** Geshani C. Bandara, Ryann D. Boudreau, William Wyatt, Steven R. Caliari

## Abstract

Biomaterial-based skeletal muscle tissue engineering approaches have largely focused on mimicking the 3D aligned architecture of native muscle, which is critical for guiding myotube formation and force transmission. In contrast, fewer studies incorporate glycosaminoglycan (GAG)-mediated biochemical cues despite their known role in regulating myogenesis and growth factor sequestration. In this study, we develop aligned collagen-GAG (CG) scaffolds using directional freeze-drying and systematically vary GAG content by incorporating GAGs of increasing sulfation levels (hyaluronic acid, chondroitin sulfate, and heparin). While all scaffold variants support myoblast adhesion, metabolic activity, and myotube alignment, heparin-modified collagen scaffolds significantly enhance myoblast metabolic activity and myogenic differentiation as measured by myosin heavy chain (MHC) expression and myotube size. We additionally show that heparin-modified scaffolds sequester and retain significantly higher levels of insulin-like growth factor-1 (IGF-1), a potent promoter of myogenesis, compared to other scaffold groups. Together, these results highlight the importance of optimizing GAG content in CG scaffolds for targeted applications and underscore the promise of heparin-modified CG scaffolds as a material platform for skeletal muscle tissue engineering.

## 1. Introduction

Skeletal muscle, comprising approximately 40% of the total body mass, consists of hierarchically organized muscle fibers embedded within a highly organized macromolecular network known as the extracellular matrix (ECM)^(1)–(5)^. While skeletal muscle possesses an intrinsic ability to regenerate following minor injury, this regenerative ability is disrupted in traumatic injuries such as volumetric muscle loss (VML)^(6)–(8)^. This is due to the removal of a large portion of muscle tissue and surrounding ECM, eliminating the structural framework and biochemical cues required for muscle cell attachment, migration, fusion into myofibers, and functional regeneration^(9)^. To address the formidable challenge of skeletal muscle tissue engineering, biomaterial-based scaffolds mimicking skeletal muscle architecture and ECM-derived biochemical signals have emerged as a promising therapeutic strategy.

The highly aligned organization of skeletal muscle is critical for generating and efficiently transferring contractile forces to enable normal locomotion^(10)^. Accordingly, recapitulating this architectural alignment has emerged as a key biomaterial design criterion for skeletal muscle tissue engineering^(10)–(14)^. For example, a previous study demonstrated that electrospun polycaprolactone/polyaniline scaffolds with aligned fibers significantly enhanced C2C12 myoblast myotube formation, as evidenced by elevated myosin heavy chain (MHC) expression^(15)^. However, layering these fibers to create 3D constructs can impede cell migration due to lower porosity and increased pore tortuosity^(16)^. Alternative methods to produce 3D constructs with aligned and interconnected pore architectures include 3D printing^(17)^ and directional freeze-drying^(18)^. A recent study by our lab demonstrated that collagen-glycosaminoglycan (CG) scaffolds with aligned micropores, produced via directional freeze-drying, promoted 3D cytoskeletal alignment and organization of C2C12 myoblasts along the collagen struts, similar to healthy skeletal muscle^(19)^.

CG scaffolds have a rich history of use as regenerative templates for tissue engineering spanning more than four decades^(20)^, with extensive investigation across a range of applications including wound healing^(21)^, peripheral nerve^(22)^, tendon^(23)^, and cartilage^(24)^ repair. Several CG scaffolds have received FDA approval, such as the Integra® Dermal Regeneration Template, which has shown success in various wound healing applications including the treatment of diabetic foot ulcers^(25)^. Chondroitin sulfate has traditionally been the most widely used glycosaminoglycan (GAG) in CG scaffolds due to its low cost and ease of sourcing when CG scaffolds were originally developed^(26),(27)^. However, CG scaffold design, specifically the selection of the GAG component, has not been optimized for skeletal muscle tissue engineering.

GAGs are linear polysaccharides composed of repeating disaccharide units that are key components of the ECM in skeletal muscle and many other tissues^(28)^. Variations in these disaccharide compositions confer different levels of sulfation, resulting in varying negative charge^(29)^. GAGs are highly polar molecules that can bind water molecules to provide hydration and mechanical support to tissues^(28)^. Their ionizable sulfate and carboxyl groups enable sequestration of proteins and cytokines such as growth factors, through electrostatic and GAG-specific interactions^(30)^. Notably, a study comparing three GAGs of increasing sulfation level (hyaluronic acid, chondroitin sulfate, and heparin) demonstrated that increased sulfation enhanced tenocyte metabolic activity within CG scaffolds^(29)^. Building on this, we hypothesized that GAG sulfation could similarly influence muscle cell proliferation and differentiation within aligned CG scaffolds. Therefore, the central goal of this work was to leverage a 3D aligned CG scaffold system to evaluate the impact of GAG sulfation on myogenesis, with the goal of establishing a material platform for skeletal muscle tissue engineering.

## 2. Materials and Methods

### 2.1 Scaffold fabrication

CG scaffolds were fabricated through directional freeze-drying of collagen-GAG suspension (**Figure 1**)^(19),(23),(32)^. A 1.5 wt/v% collagen suspension was made by mixing type I collagen derived from bovine Achilles tendon (Sigma Aldrich) in 0.05 M acetic acid using a high shear homogenizer while maintaining the temperature at 4°C via a recirculating chiller to prevent collagen denaturation. Precooled GAG solutions consisting of hyaluronic acid (HA, 19 kDa, Lifecore), chondroitin sulfate derived from bovine cartilage (CS, 10-40 kDa, Sigma Aldrich), or heparin derived from porcine intestinal mucosa (HP, 5-15 kDa, Sigma Aldrich) at 0.1 wt/v% were made by vortexing GAG in 0.05 M acetic acid and added dropwise to the collagen suspension under homogenization. The homogenized CG suspension was then pipetted into a thermally mismatched Teflon mold with a copper base, facilitating longitudinal heat transfer.

**Figure 1.**
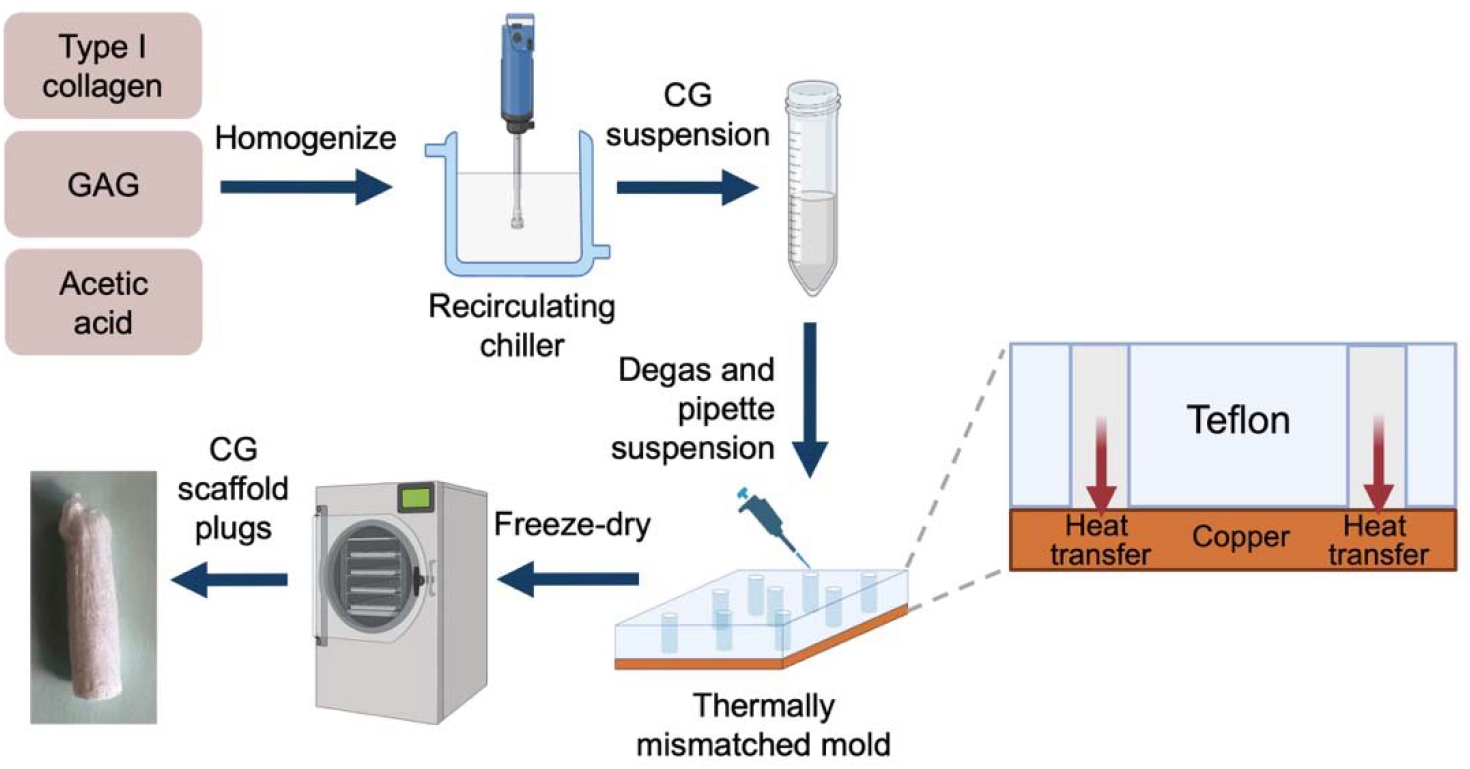
Schematic illustration of the collagen-glycosaminoglycan (CG) scaffold fabrication process. The suspension was pipetted into a thermally mismatched mold and placed in a pre-cooled freeze-dryer shelf at −10 °C. During the initial freezing phase, ice crystals formed longitudinally due to unidirectional heat transfer through the copper bottom plate. These ice crystals subsequently sublimated during the vacuum phase of freeze-drying, leaving behind scaffolds with longitudinally aligned interconnected pore structure.

### 2.2 Scaffold hydration, crosslinking, fluorescent labeling, and sterilizing

Following freeze-drying, the scaffolds underwent dehydrothermal crosslinking under vacuum at 105°C for 24 h. Scaffold plugs (6 mm diameter, 2.5-3 mm height, cut in half) were immersed in 70% ethanol for initial hydration followed by rinsing with PBS. Amines on the scaffold collagen backbone were labeled with fluorophore by gently shaking scaffold plugs in a 2 μM solution of Alexa Fluor 568 NHS ester in PBS for 20 min. Finally, the scaffolds were chemically crosslinked using 1-ethyl-3-(3-dimethylaminopropyl) carbodiimide hydrochloride (EDC) and N-hydroxysulfosuccinimide (NHS) at a molar ratio of 5:2:1 EDC:NHS:COOH, where COOH represents the carboxylic acid content of the collagen^(33)^. EDC/NHS activates carboxylic acid groups to covalently crosslink with primary amines in collagen, improving scaffold mechanical stability. Additionally, this crosslinking method increases CG scaffold resistance to biodegradation via enzymes such as collagenase and chondroitinase^(34)^. Following chemical crosslinking, the scaffolds were sterilized by immersion in 70% ethanol for 2 h, then transferred to sterile PBS and incubated overnight before cell seeding.

### 2.3 Scanning electron microscopy analysis

Dry scaffold pore structure was analyzed using scanning electron microscopy (SEM). A backscatter electron detector (BSD) at 60 Pa of a Phenom ESEM at 10 kV was used to capture images. Pore anisotropy was characterized using ImageJ OrientationJ plugin in longitudinal (parallel to freezing direction) and transverse (perpendicular to freezing direction) plane images.

### 2.4 Cell culture

Immortalized mouse myoblasts (C2C12, ATCC) were utilized at passages 2-6. Cells were cultured in standard T75 tissue culture flasks using growth media consisting of high-glucose Dulbecco’s Modified Eagle’s Medium (DMEM) supplemented with 10 v/v% fetal bovine serum (FBS, Gibco) and 1 v/v% antibiotic-antimycotic solution (Invitrogen). Cells were maintained until 80% confluency prior to passaging or experimental use. For differentiation, cells were switched to differentiation media composed of DMEM supplemented with 2 v/v% horse serum (Gibco) and 1 v/v% antibiotic-antimycotic (Invitrogen). All cultures were maintained at 37□°C and 5% CO_2_.

### 2.5 Scaffold culture conditions

Hydrated, crosslinked, and sterilized scaffolds were rinsed with sterile PBS and pre-incubated in growth media for 30 min prior to cell seeding. Three or four scaffolds per group were then transferred into a well of an ultralow attachment 6-well plate, containing 1 mL/scaffold of cell suspension with 3 × 10^5^ cells/mL. Plates were placed on an orbital shaker and agitated at 100 rpm overnight at 37□°C in a humidified atmosphere with 5% CO_2_ to promote uniform cell attachment throughout the 3D scaffolds. Media was changed the next day and subsequently changed every other day. Scaffolds were maintained in growth media for 4 days before switching to differentiation media to induce myogenic differentiation for an additional 4 or 7 days.

### 2.6 Cell metabolic activity

Cell metabolic activity was assessed non-destructively using the alamarBlue assay^(23)^. Cell-seeded scaffolds were incubated in a 10% alamarBlue solution for 70 min on a shaker at 100 rpm. During incubation, viable cells facilitate mitochondrial and cytoplasmic redox reactions, reducing the non-fluorescent blue resazurin present in the alamarBlue solution to fluorescent pink resorufin. The resulting fluorescence signal was quantified using a plate reader (Tecan M200).

### 2.7 Immunocytochemistry and confocal imaging

At the end of the culture period, scaffolds were fixed in 10% neutral buffered formalin. Following fixation, samples were permeabilized in 0.1% Triton X-100 in PBS for 30 min and then blocked in 3 w/v% bovine serum albumin (BSA) in PBS for 1 h to reduce nonspecific antibody binding. Scaffolds were subsequently incubated overnight at 4□°C with a primary anti-myosin heavy chain (MHC) antibody (Myosin 4 monoclonal antibody (MF20), eBioscience) diluted 1:200 in 3 w/v% BSA. After incubation, scaffolds were rinsed three times with PBS and then incubated for 2 h at room temperature with Alexa Fluor 488-conjugated goat anti-mouse IgG (1:200 dilution in PBS; Thermo Fisher Scientific) as the secondary antibody. Following secondary antibody labeling, scaffolds were washed three times with 0.05 v/v% Tween in PBS, then incubated with DAPI (4′,6-diamidino-2-phenylindole) for 5 min for nuclear staining. Finally, samples were rinsed twice with PBS and stored in the dark at 4□°C until imaging.

Imaging was performed using a Cytation C10 confocal microscope. The collagen backbone, MHC expression, and nuclei were visualized using TRITC, GFP, and DAPI channels, respectively. Z-stack images (100 μm thickness) were acquired from five fields of view per scaffold in the longitudinal plane. Maximum intensity projections of the z-stacks were used for analysis. Cells were segmented using both nuclei and MHC channels using Omnipose^(35)^. Segmentation masks were manually corrected, and cell morphological properties were quantified using the region_props function from the scikit-image Python package. Myotube formation and maturation were assessed based on MHC expression, and fusion index was calculated as the percentage of total nuclei in myotubes by overlaying the MHC and DAPI channels.

### 2.8 Growth factor (Insulin-like growth factor 1 (IGF-1) and basic fibroblast growth factor (bFGF)) pulldown sequestration and retention assay

To quantify the extent of IGF-1 and bFGF sequestration, a pulldown assay was performed, with IGF-1/bFGF levels measured using enzyme-linked immunosorbent assay (ELISA) kits. For the assay, five hydrated and crosslinked scaffolds per group were incubated overnight at 37 °C in a single well of an ultralow attachment 6-well plate. Each well contained 4 mL of PBS solution with 100 ng/mL IGF-1 (Prospec-CYT-229) or bFGF (ProSpec, CYT-386) and 1 w/v% BSA. A control group, consisting of wells containing IGF-1/bFGF solution without scaffolds, was included to account for non-specific retention. Following incubation, the remaining growth factor concentration in each well was quantified using an ELISA kit (R&D Systems) with samples diluted 1:49 in PBS according to the manufacturers protocol. Relative growth factor pulldown (%) was calculated by determining the difference between IGF-1/bFGF levels in the experimental and control groups, normalized to the control group.

IGF-1 and bFGF retention within scaffolds during cell culture was quantified at three time points: day 4 in growth media, day 8 (4 days in differentiation media), and 11 (7 days in differentiation media). At each time point scaffolds were rinsed with cold PBS to remove serum proteins or residual media, then lysed in 1 mL of RIPA lysis and extraction buffer (1 mL/scaffold, Thermo Scientific,) supplemented with Halt™ protease inhibitor cocktail (10 µL per 1 mL, Thermo Scientific) for 30 min on ice. Scaffolds were placed individually into low protein-binding microcentrifuge tubes and vortexed every 10 min to enhance lysis. Lysates were collected, and scaffolds were transferred to fresh low protein-binding tubes with 1.5 mL of collagenase I solution (3 mg/mL, Sigma Aldrich) and digested for 72 h at 37□°C. After completion of digestion, samples were centrifuged at 12000 g for 15 min, and supernatants were collected. Mouse and rat IGF-1 (MG100) and Mouse and rat bFGF (MFB00) ELISA kits were used for IGF-1 and bFGF experiments respectively. All samples and standards were run in duplicate and fluorescence intensity was measured at 450 and 570 nm using a Tecan M200 plate reader.

### 2.9 Statistical analysis

Statistical tests were performed using one-way or two-way ANOVA followed by Tukey’s HSD post hoc test using GraphPad Prism 10. Statistically significant differences are indicated by *, **, ***, and **** corresponding to *p* < 0.05, 0.01, 0.001, or 0.0001 respectively. Heights of bar graphs correspond to the mean with standard deviation error bars and scatter plots are overlaying the bars indicating individual data points. Each experiment involved *n* = 3-6 scaffolds per experimental group while immunocytochemical analyses were done on five ROIs per scaffold.

## 3 Results and Discussion

### 3.1 Varying GAG type does not alter CG scaffold pore structure

Previous work from our team demonstrated successful fabrication of aligned porous CG scaffolds incorporating CS^(19),(23),(32)^. To evaluate the influence of GAG type (HA, CS, or HP) on CG scaffold pore structure, SEM imaging was performed (**Figure 2**). In the longitudinal plane, all three scaffold groups exhibited pore anisotropy in the direction of heat transfer, whereas the transverse plane images revealed more randomized pore orientation. This observation was quantitatively supported by quantification of the distribution of pore orientation angles in the polar plots, which showed a concentration of alignment frequencies in the longitudinal plane around 0°, indicating vertical alignment, and a more dispersed distribution in the transverse plane. These observations confirm the successful fabrication of aligned porous scaffolds regardless of the GAG type incorporated.

**Figure 2:**
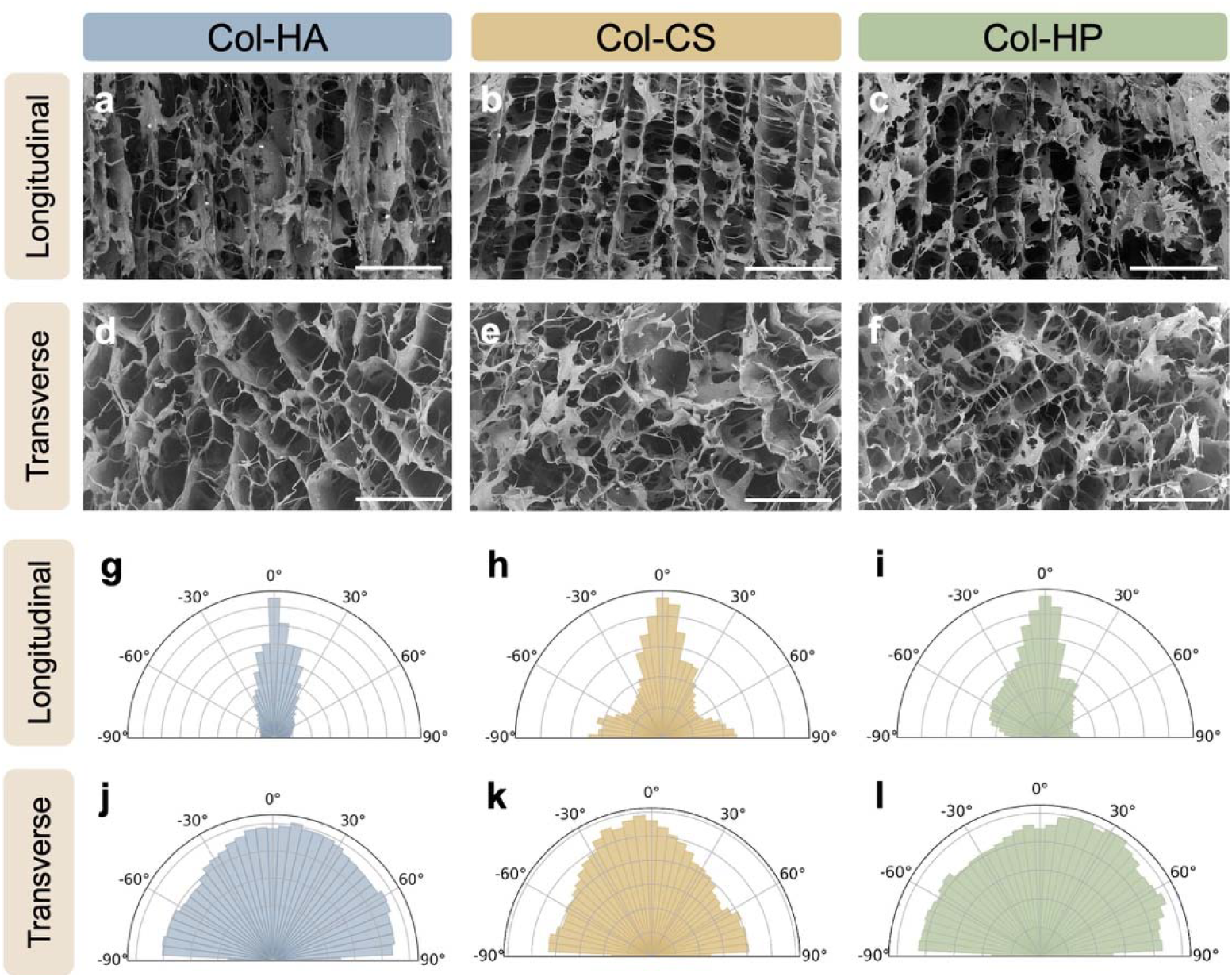
Scaffolds showed longitudinally aligned pores independent of GAG type. a-c) SEM images show aligned pores in the longitudinal plane mimicking native skeletal muscle tissue and (d-f) more randomized pores in the transverse plane. Pore orientation was quantified using polar plots for the (g–i) longitudinal and (j–l) transverse planes. Scale bars: 300 μm; *N* = 3 scaffolds per group.

### 3.2 Heparin-modified scaffolds significantly enhance myoblast metabolic activity

Previous studies demonstrated that CS-modified CG scaffolds support sustained myoblast metabolic activity^(19),(32)^. To evaluate and compare cellular behavior in scaffolds as a function of GAG type, scaffolds were seeded with myoblasts and cultured in myoblast growth media for 4 days, followed by differentiation media for 7 days (**Figure 3**). Cell metabolic activity was then assessed non-destructively using the alamarBlue cell metabolic activity assay. Globally, the scaffolds supported increased cell metabolic activity over the 11 day culture period. Notably, HP-incorporated scaffolds supported significantly higher cell metabolic activity compared to HA- and CS-incorporated scaffolds by day 6. While HA scaffolds showed a statistically significant improvement over CS scaffolds on days 1 and 4, this effect was less prominent than the enhanced activity observed with HP by the end of the culture period. Additionally, a slight decrease in the metabolic activity was observed on day 11 compared to day 8 across all groups. This reduction may reflect the transition of cells from a more proliferative to differentiated state as they exit the cell cycle and become less metabolically active.

**Figure 3:**
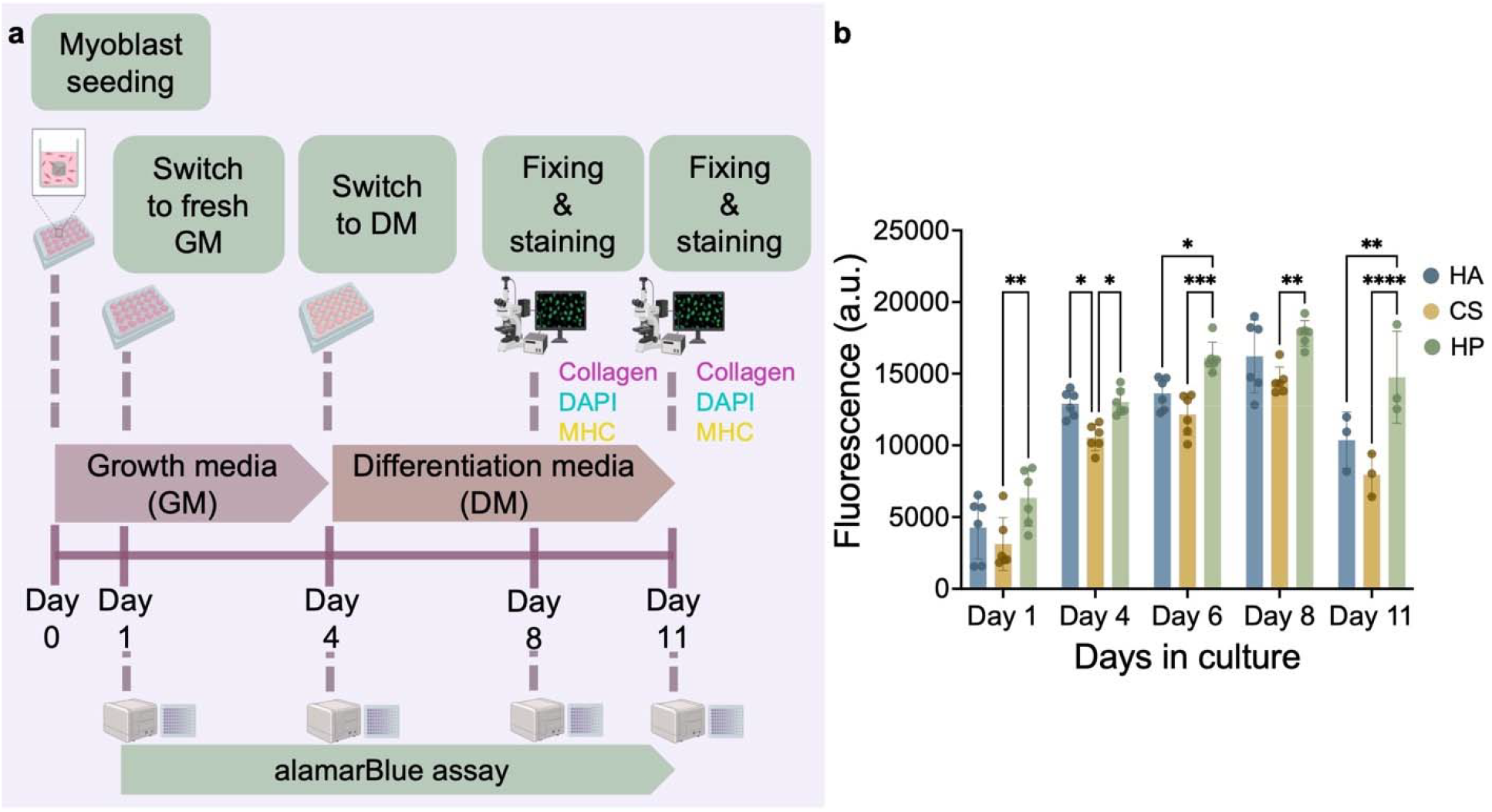
CG scaffolds supported sustained myoblast metabolic activity. a) Schematic illustrating CG scaffold cell seeding and culture timeline. b) Fluorescence values obtained from an alamarBlue cell metabolic activity assay showed significantly increased cell metabolic activity in HP-incorporated scaffolds over the 11-day culture period. Two-way ANOVA with Tukey’s HSD post hoc tests. *: p < 0.05, **: p < 0.01, ***: p < 0.001, ****: p < 0.0001. *N* = 3-6 scaffolds per experimental group.

A similar trend was observed in a previous study comparing HA-, CS-, and HP-incorporated collagen scaffolds^(31)^. In that work, HP-incorporated scaffolds supported significantly improved tenocyte and human mesenchymal stromal cell metabolic activity compared to HA and CS scaffolds. The authors correlate this higher metabolic activity to the greater retention and sequestration of insulin-like growth factor-1 (IGF-1) within HP scaffolds, emphasizing that higher sulfation facilitates stronger growth factor binding, sustained local signaling, and enhanced cell metabolic activity. Consistent with these findings, another study demonstrated increased endothelial cell metabolic activity in composite silk-based vascular scaffolds modified with heparin compared to unmodified scaffolds^(36)^. Similarly, another study highlighted improved myoblast metabolic activity in heparinized decellularized skeletal muscle ECM scaffolds treated with platelet rich plasma, underscoring the role of scaffold sulfation in regulating scaffold performance by sequestering growth factors^(37)^. Taken together, these results motivated us to further evaluate the capacity of CG scaffolds for supporting myogenic differentiation as well as growth factor sequestration and retention.

### 3.3 Myotubes aligned along scaffold pore orientation induced by directional freeze-drying, independent of GAG type

After confirming scaffold structural alignment and sustained myoblast metabolic activity, myoblast organization and differentiation into myotubes within CG scaffolds incorporating HA, CS, or HP was evaluated by immunocytochemical analysis of myosin heavy chain (MHC), a muscle cell differentiation marker, using confocal microscopy after 4 and 7 days in differentiation media. Comparable myotube alignment was observed across all three scaffold groups on day 4 (**Figure S1**). Regardless of GAG type, all scaffolds largely maintained myotube alignment ay day 7. The strong overlap between myotube alignment and scaffold pore alignment further indicates that the contact guidance provided by the aligned CG scaffold backbone supported myotube organization (**Figure 4**).

**Figure 4:**
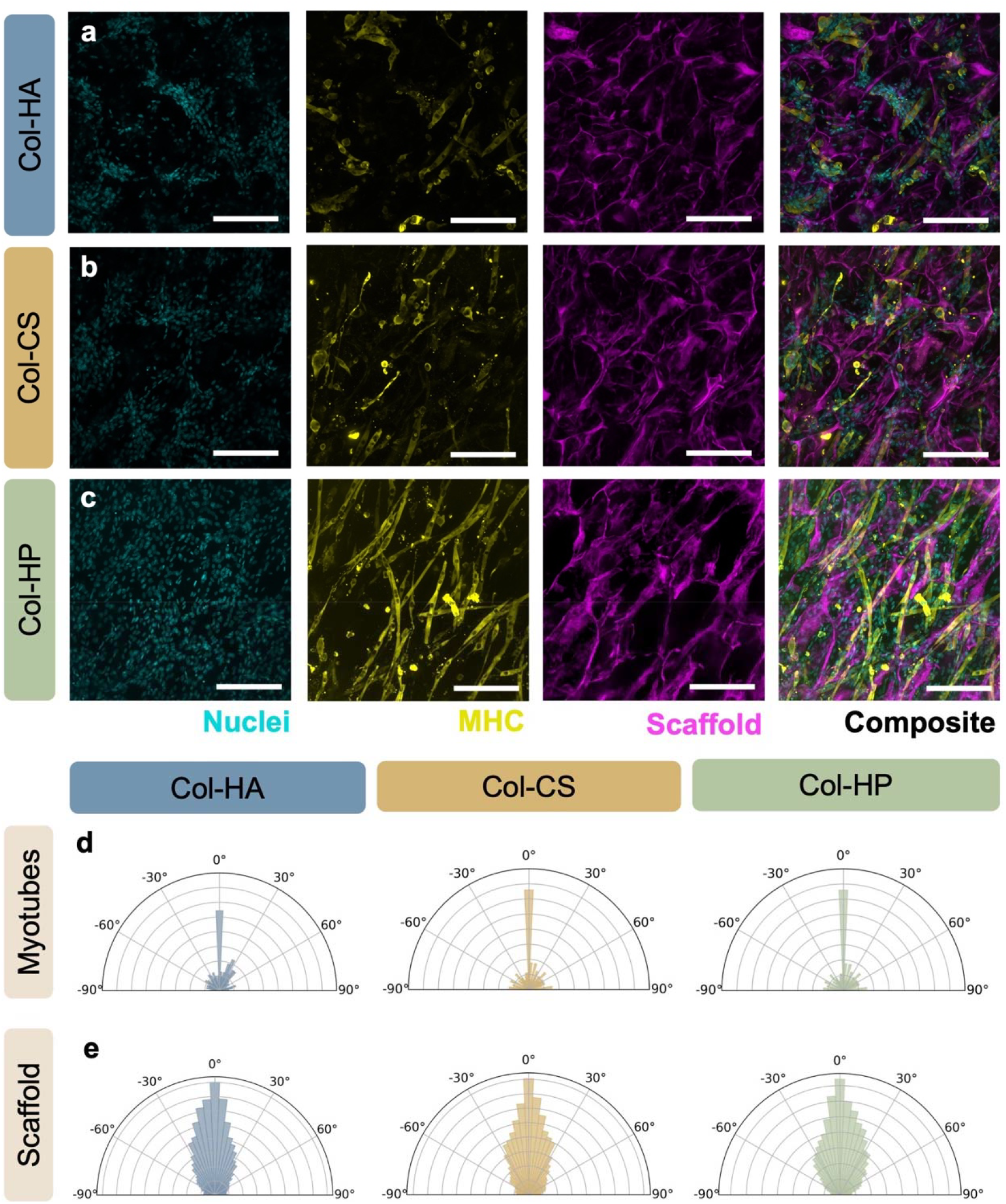
Aligned scaffolds promoted comparable myotube organization after 7 days in differentiation media independent of GAG type. a) Confocal microscopic images of myotubes in collagen-HA (Col-HA), b) collagen-chondroitin sulfate (Col-CS), and c) collagen-heparin (Col-HP) scaffolds reveal similar levels of myotube alignment along the oriented pore structure. d) Corresponding polar plots quantify myotube orientation normalized to total myotubes count, confirming consistent alignment with e) scaffold backbone orientation across all scaffold groups. Scale bars: 200 μm; *N* = 3 scaffolds per experimental group.

### 3.4 Heparin-modified scaffolds supported sustained myogenesis

Following confirmation of myotube alignment and organization in 3D aligned CG scaffolds, independent of GAG type, we next quantified myogenesis via analysis of MHC staining density, myotube size, and fusion index. Similar levels of myogenic differentiation were quantified across all three CG scaffold groups following 4 days in differentiation media (**Figure S2**). However, by day 7 in differentiation media, HP-incorporated scaffolds exhibited significantly improved *in vitro* myogenesis compared to HA and CS groups as measured by MHC+ area, with significantly higher myotube length and width compared to the CS group (**Figure 5**). These results highlight that HP scaffolds supported the formation and maintenance of myotubes over time, whereas myotube presence did not persist as much in the other two groups.

**Figure 5:**
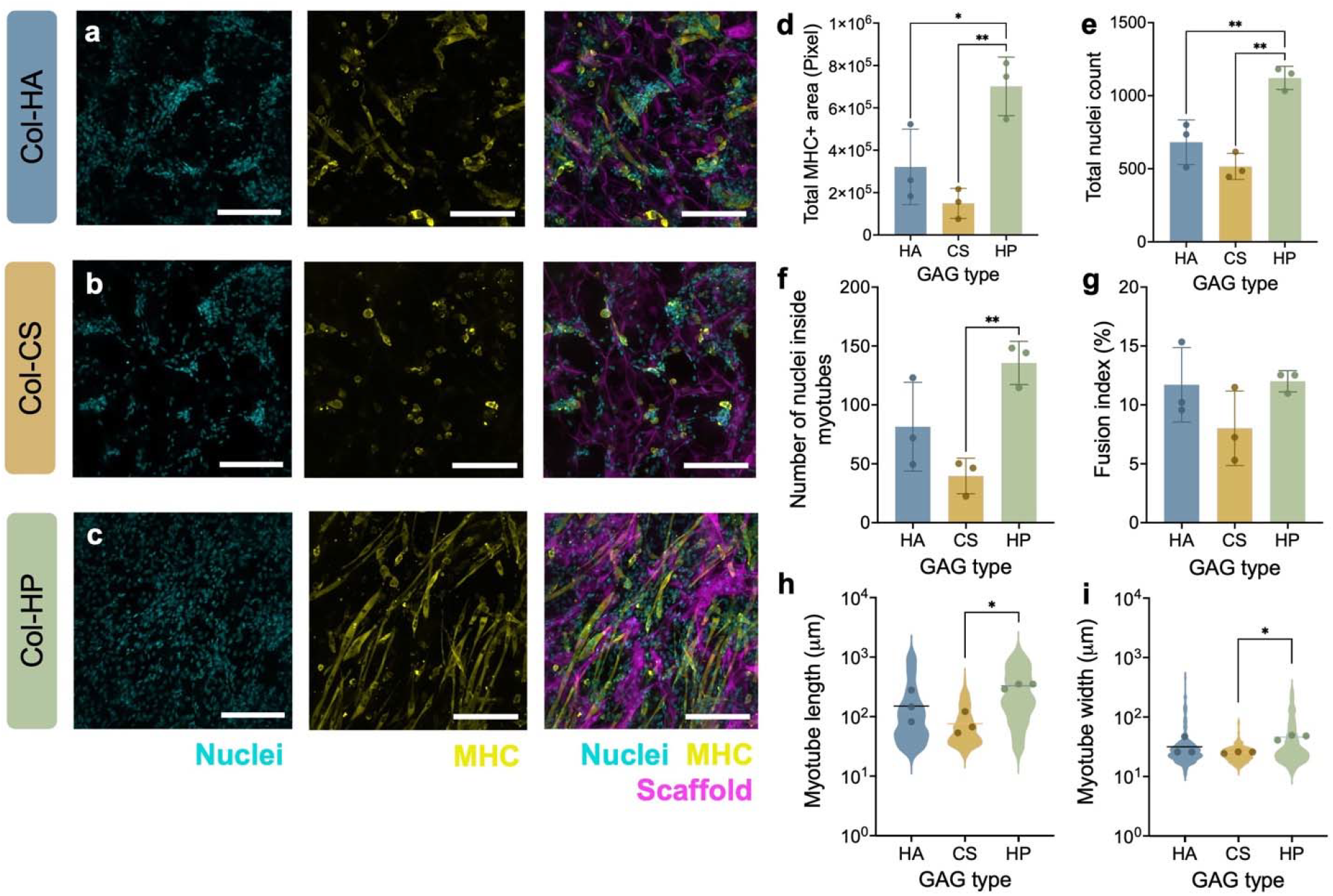
Heparin-modified scaffolds supported significantly enhanced and sustained myogenic differentiation after 7 days in differentiation media. a) Confocal microscopic images of myotubes in collagen-HA (Col-HA), b) collagen-chondroitin sulfate (Col-CS), and c) collagen-heparin (Col-HP) scaffolds. d) Analysis of myogenic differentiation in CG scaffolds containing HA, CS, or HP revealed significantly higher total MHC+ area, e) total nuclei count, and f) number of nuclei inside myotubes in HP-modified scaffolds. g) Fusion index (percentage of nuclei inside myotubes) was similar across experimental groups. h) myotube length and i) width were significantly increased in HP-modified scaffolds compared to CS-modified scaffolds. One-way ANOVA with Tukey’s HSD post hoc tests. *: p < 0.05, **: p < 0.01. Scale bars: 200 μm; *N* = 3 scaffolds per experimental group.

Although the fusion index was similar across all three groups, HP scaffolds showed higher total nuclei counts as well as a greater number of nuclei inside myotubes. Further analysis of myotube fractions revealed a significantly lower fraction of immature myotubes with just two nuclei in the HP-incorporated scaffolds compared to CS-incorporated scaffolds (**Figure S3**). These findings indicate that HP scaffolds not only promoted myotube formation, but also supported their maturation.

### 3.5 Heparin-modified scaffolds support increased growth factor sequestration

We hypothesized that the sustained myogenic differentiation observed in HP scaffolds was due to enhanced growth factor (GF) sequestration and retention over time. To test this hypothesis, GF sequestration and retention assays were performed on scaffolds at various culture time points. First, a modified GF pulldown assay was used to quantify and compare the GAG-mediated GF sequestration across the three scaffold groups prior to cell seeding (**Figure S4**). Two representative GFs were selected: basic fibroblast growth factor (bFGF), known to promote muscle cell proliferation^(38)^, and insulin-like growth factor (IGF-1) which promotes both muscle cell proliferation and differentiation^(39)–(43)^. HP-incorporated scaffolds exhibited significantly increased bFGF sequestration compared to CS-incorporated scaffolds, and significantly enhanced IGF-1 pulldown compared to both HA- and CS-incorporated scaffolds. The primary mechanism of GF sequestration, retention, and stabilization is charge-based interactions between the negatively-charged sulfate and carboxylic groups of GAGs and positively-charged domains of GFs^(44),(45)^. Among these GAGs, HP possesses the highest degree of sulfation resulting in a higher negative charge compared to HA and CS, conferring strong affinity for positively charged GFs. Previous studies indicated that high hydration and the porous structure of HA-based biomaterials can promote physical adsorption and entrapment of GFs^(46),(47)^. Accordingly, this may explain the slight increase in GF sequestration observed in HA scaffolds compared to CS scaffolds, despite the increased sulfation levels in CS scaffolds.

To quantify the amount of GFs retained inside the scaffolds during cell culture, cultured scaffolds were lysed and digested, and the amounts of bFGF and IGF-1 were measured using ELISA. While all scaffold groups retained similar bFGF levels, HP-incorporated scaffolds exhibited significantly higher IGF-1 levels at all time points evaluated (days 4, 8, and 11) compared to other groups (**Figure 6, Figure S5**). IGF-1 levels increased over the culture period across all groups, with the most pronounced accumulation observed in the HP scaffolds. Given that IGF-1 is a primary driver of myoblast differentiation, this enhanced retention is expected to facilitate myogenic differentiation within scaffolds.

**Figure 6:**
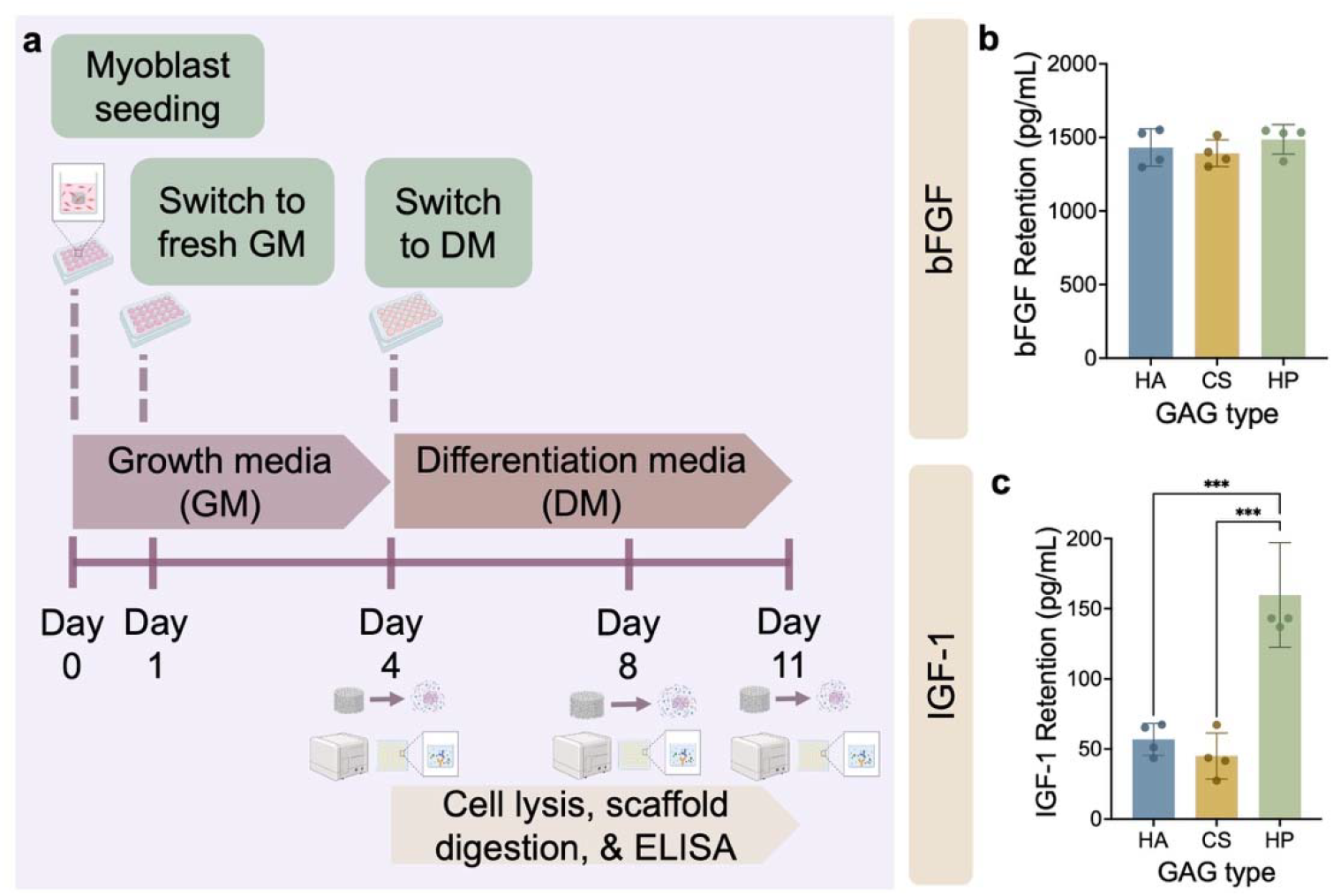
Increased GAG sulfation enhances scaffold IGF-1 retention. a) Schematic of the growth factor retention quantification using ELISA. b) While relatively minor differences in bFGF levels were quantified after 11 days of culture, c) significantly higher levels of IGF-1 were found in HP-modified scaffolds compared to HA- and CS-modified groups. One-way ANOVA with Tukey’s HSD post hoc tests. ***: p < 0.001. *N* = 4 scaffolds per experimental group.

## 4 Conclusions

In this study, we investigated the role of GAG type on *in vitro* myogenesis in 3D structurally aligned CG scaffolds by incorporating HA, CS, and HP representing increasing sulfation levels. Our findings demonstrated that highly sulfated HP-incorporated scaffolds significantly enhanced myoblast metabolic activity and supported the formation and maintenance of aligned multinucleated myotubes compared to less sulfated GAGs. To explore the underlying mechanism, we showed that HP-modified scaffolds promoted markedly greater sequestration and retention of the myogenic growth factor IGF-1 compared to HA- and CS-modified scaffolds. Together, these results highlight the importance of optimizing GAG content in CG scaffolds for targeted applications and underscore the promise of HP-incorporated CG scaffolds as a material platform for skeletal muscle tissue engineering.

## Acknowledgments

The authors thank James Gentry for developing the Omnipose code used to analyze myotube confocal images. SEM images were acquired at the University of Virginia Nanoscale Materials Characterization Facility (NMCF). This work was supported by the NIH (R01AR078866). The content is solely the responsibility of the authors and does not necessarily represent the official views of the National Institutes of Health.

## Data availability statement

Key data supporting this article have been included as part of the Supplementary Information.

