## Supplementary Information for "Heparin-modified aligned collagen scaffolds enhance *in vitro* myogenesis"

### Supplemental Information

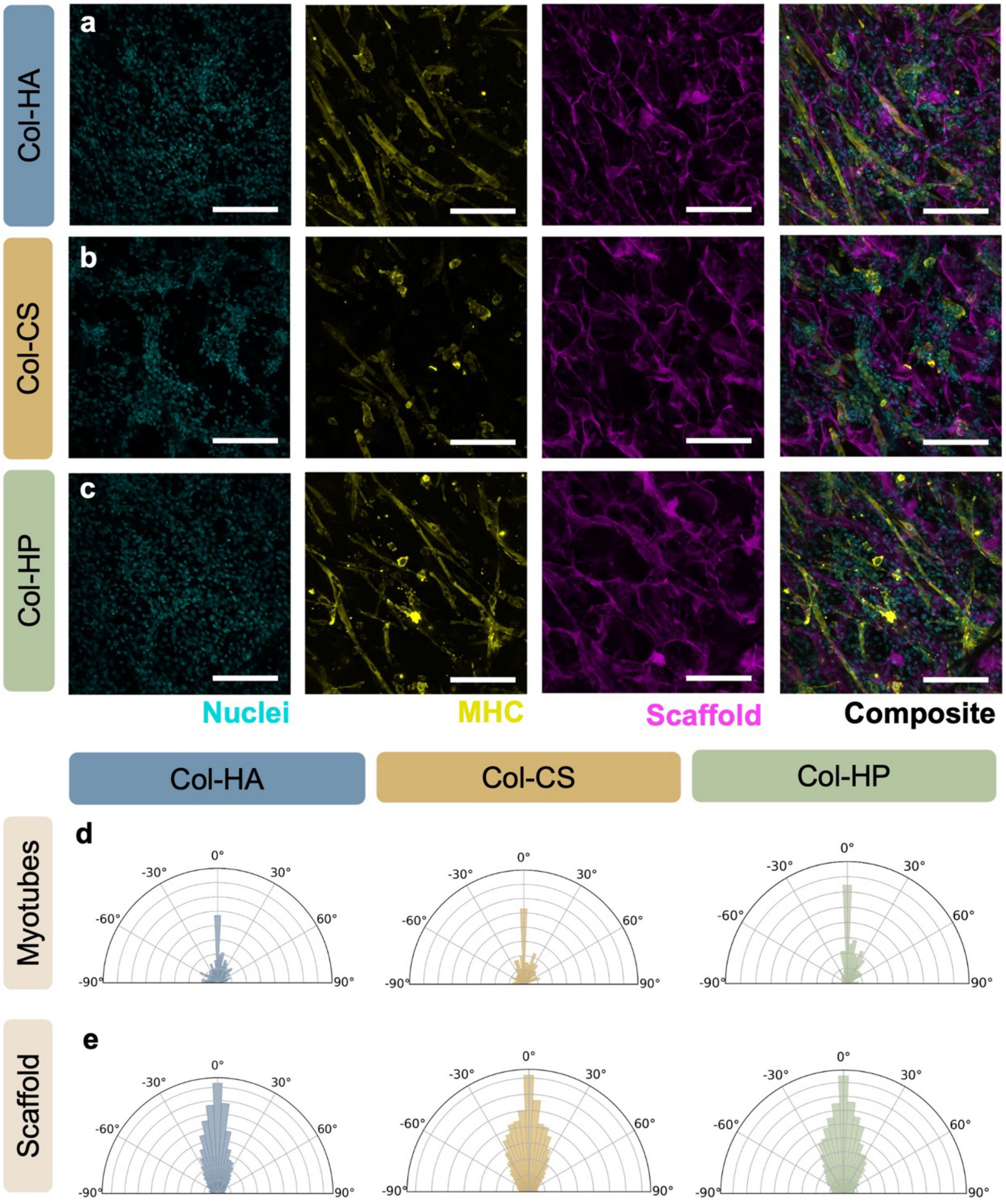


**Figure S1: Aligned scaffolds promoted comparable myotube organization after 4 days in differentiation media independent of GAG type.** a) Confocal microscopic images of myotubes in collagen-HA (Col-HA), b) collagen- chondroitin sulfate (Col-CS), and c) collagen-heparin (Col-HP) scaffolds reveal similar levels of myotube alignment along the oriented pore structure. d) Corresponding polar plots quantify myotube orientation normalized to total myotubes count, confirming consistent alignment with e) scaffold backbone orientation across all scaffold groups. Scale bars: 200 µm; *N* = 3 scaffolds per experimental group.

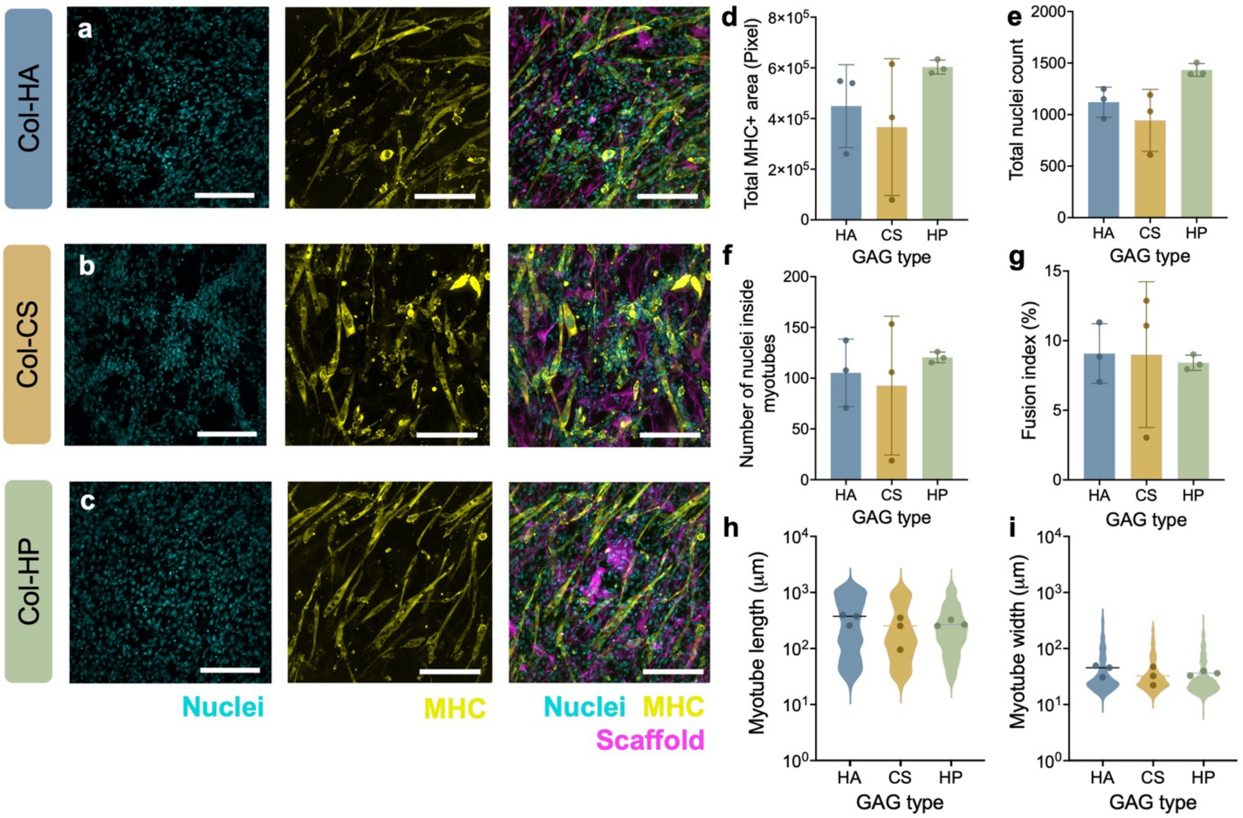


**Figure S2: Scaffolds supported similar myogenic differentiation after 4 days in differentiation media.** a) Confocal microscopic images of myotubes in collagen-HA (Col-HA), b) collagen-chondroitin sulfate (Col-CS), and c) collagen-heparin (Col-HP) scaffolds. d) Analysis of myogenic differentiation in CG scaffolds containing HA, CS, or HP revealed similar total MHC+ area, e) total nuclei count, f) number of nuclei inside myotubes, g) fusion index (percentage of nuclei inside myotubes), h) myotube length, and i) myotube width. One-way ANOVA with Tukey's HSD post hoc tests. Scale bars: 200 μm; *N* = 3 scaffolds per experimental group.

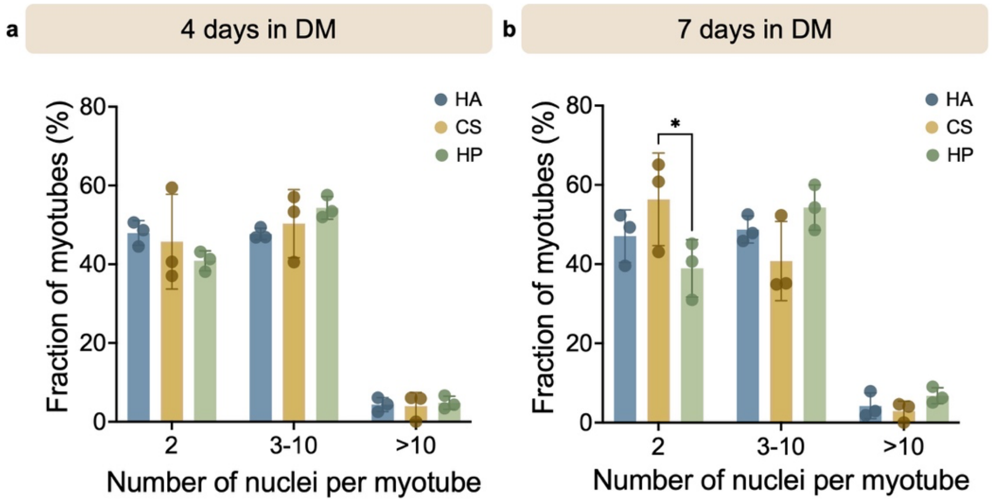


**Figure S3: Heparin-modified scaffolds show lower fraction of immature myotubes after 7 days in differentiation media.** a) Analysis of myogenic differentiation in CG scaffolds containing HA, CS, or HP revealed distributions of number of nuclei in myotubes after 4 days in differentiation media. b) By day 7 in differentiation media, HP-incorporated scaffolds had a significantly lower fraction of immature myotubes with just two nuclei compared to the CS-incorporated scaffolds. Two-way ANOVA with Tukey's HSD post hoc tests. ✻: $p < 0.05$. $N = 3$ scaffolds per experimental group.

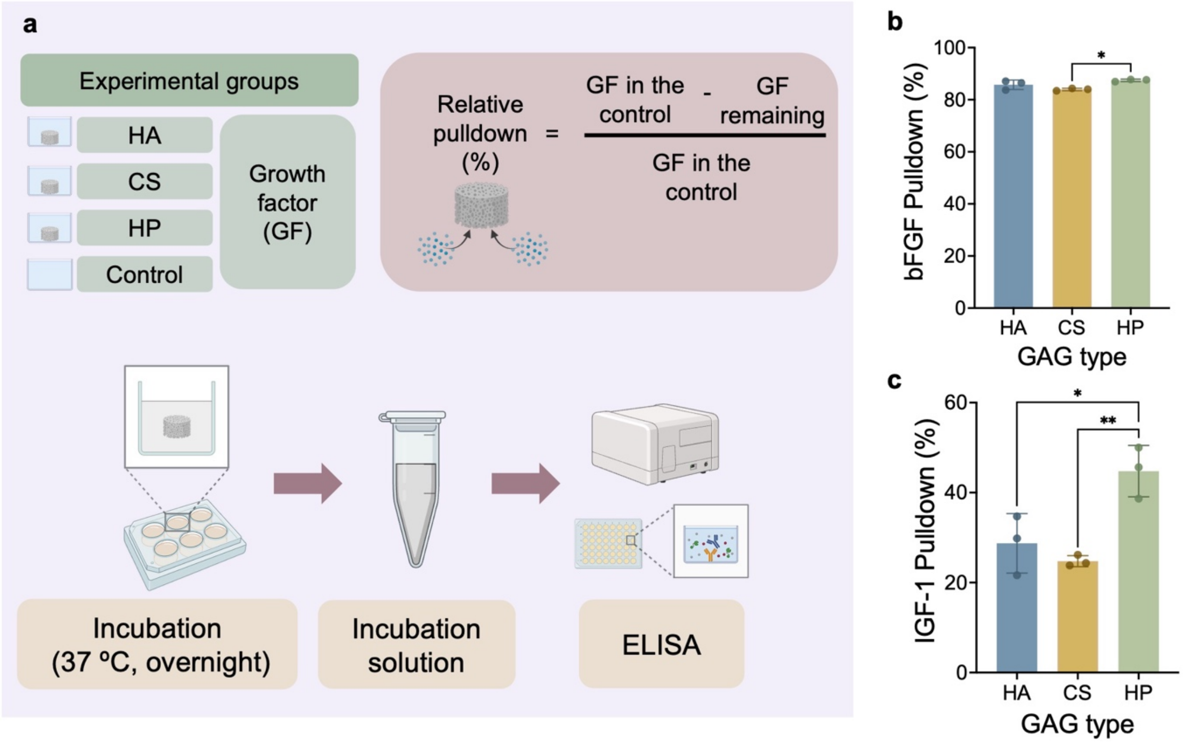


**Figure S4: Increased GAG sulfation enhances scaffold growth factor sequestration.** a) Schematic of the growth factor (GF) pulldown assay where bFGF and IGF-1 levels are quantified by ELISA. b) HP-incorporated scaffolds showed slightly higher bFGF pulldown compared to HA- and CS-modified scaffolds. c) HP-incorporated scaffolds demonstrated significantly higher relative IGF-1 pulldown. One-way ANOVA with Tukey's HSD post hoc tests. ✻: $p < 0.05$, ✻✻: $p < 0.01$. $N = 3$ scaffolds per experimental group.

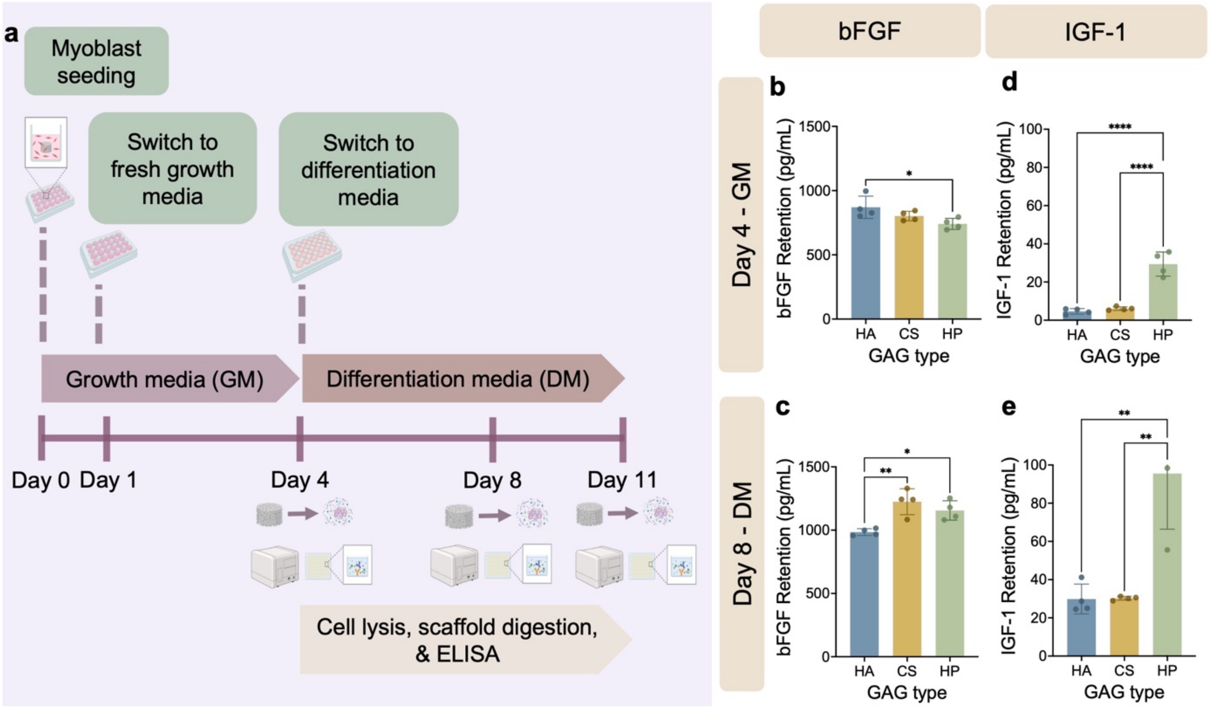


**Figure S5: Increased GAG sulfation enhances scaffold growth factor retention at intermediate culture time points.** a) Schematic of the growth factor retention quantification using ELISA. b) While relatively minor differences in bFGF levels were quantified at days 4 and c) 8 of culture, d) significantly higher levels of IGF-1 were found in HP-modified scaffolds after both day 4 and e) day 8 of culture. One-way ANOVA with Tukey's HSD post hoc tests. ✻: $p < 0.05$, ✻✻: $p < 0.01$, ✻✻✻: $p < 0.001$, ✻✻✻✻: $p < 0.0001$. $N = 4$ scaffolds per experimental group.
